# ‘Erythritol’, a safe natural sweetener exhibits multi-stage anti-malarial activity by blocking *Plasmodium falciparum* membrane transporter ‘aquaporin’

**DOI:** 10.1101/2022.04.28.489976

**Authors:** Jyoti Kumari, Vikash Kumar, Ankita Behl, Raj Kumar Sah, Swati Garg, Soumya Pati, Kirandeep Samby, Jeremy Burrows, Narla Mohandas, Shailja Singh

## Abstract

The increased resistance of human malaria parasite *Plasmodium falciparum* to currently used drugs necessities the development of novel anti-malarials. In the present study, we examine the potential of erythritol, a sugar substitute, for therapeutic intervention that target a multifunctional transporter protein *Plasmodium* aquaglyceroporin (*Pf*AQP) responsible for maintaining hydro-homeostasis. We show that erythritol effectively inhibited growth and progression of asexual blood stage malaria parasite by suppressing invasion and egress processes. It inhibited the liver stage (sporozoites) and transmission stage parasite (gametocytes) development that suggest its multi-stage, transmission-blocking potential. Interestingly, erythritol inhibited *in vivo* growth of malaria parasite in mouse experimental model. It was more effective in inhibiting parasite growth both *in vivo* and *in vitro* when tested together with a known anti-malarial ‘artesunate’. No Evans blue staining in treated mice indicated erythritol mediated protection of blood–brain barrier integrity in mice infected with *P. berghei*. Additionally, erythritol showed cytokine-modulating effect which suggest its direct effect on the host immune system. Our results of cellular thermal shift assay and ammonia detection assay demonstrate that erythritol binds with *Pf*AQP and reduce the amount of ammonia release across the parasite respectively. We performed functional complementation assays which suggest that *Pf*AQP expression in yeast mutant restores its growth in hyperosmotic conditions but showed reduced growth in the presence of erythritol, suggesting erythritol as an inhibitor of *Pf*AQP. Overall, our data bestow erythritol as a promising new lead compound with an attractive antimalarial profile and could possibly be combined with known drugs without losing its efficacy.

## Introduction

*Plasmodium falciparum* (*Pf*) is the major cause of malaria that is responsible for ∼0.4 million deaths annually. Despite various efforts to reduce the global burden of the disease, malaria remains a major killer of children under 5 years of age. Prophylaxis and chemotherapeutics are the current approaches used to combat malaria. However, emergence of resistant *Plasmodium* strains impedes the efforts and strategies for control and eradication of this disease. Artemisinin, an extremely potent and fast-acting anti-malarial drug was proposed to have indispensable role in malaria therapies when used in combination with other anti-malarials such as mefloquine or lumenfantine [1]. However, resistance to artemisinin combination therapy has also been reported which further threatens to compromise the global endeavours to control this disease [2, 3]. Strategies adopted for effective disease control should aim to identify new targets and their anti-malarials to overcome the impact of this deadly disease. One such plausible target in *Plasmodium falciparum* is Aquaporin.

The aquaporins belong to a family of small (24-30 kDa) pore-forming integral membrane proteins that act as membrane transporters and play significant role in maintaining water homeostasis [4, 5, 6]. They are widely distributed throughout nature in diverse taxa including archaea, eubacteria, fungi, plants and animals [6]. Aquaporins are classified into two subgroups namely aquaporins and aquaglyceroporins. The classical aquaporins are involved in transporting mainly water whereas the aquaglyceroporins conduct glycerol and various other small, uncharged molecules such as polyols, urea and arsenic [6]. Structurally aquaporins are conserved proteins. They assemble to form homotetramers in membranes where each monomer forms a single pore and function independently [6, 7]. Aquaporin monomer consists of six transmembrane helical segments and two short helices with cytoplasmically oriented amino and carboxy termini. Sequence alignment of the aquaporins show several highly conserved motifs, including “NPA” motifs, and “AEFL” and “HW[V/I][F/Y]WXGP” sequences [6,7].

Genome of *Plasmodium falciparum* encodes for two aquaporins [8]. One is the classical aquaporin (PF3D7_0810400) and other aquaglyceroporin (*Pf*AQP; PF3D7_1132800). Several reports define the structure and function of *Pf*AQP while classical aquaporin remains a putative protein with no existing information. Hansen et al. reported that *Pf*AQP is expressed during the parasite asexual blood stages namely rings, trophozoites and schizonts, and is localized solely to the parasite [9]. *Pf*AQP shares highest similarity to the *Escherichia coli* glycerol facilitator (50.4%), but both canonical NPA motifs in the pore region are replaced with NLA and NPS, respectively [9]. Several reports suggest multifunctional feature of *Pf*AQP owing to its permeability for water, urea, small polyols and carbonyl compounds [9-11]. Besides, *Pf*AQP was found to be permeable for glycerol, suggesting its role in providing access to serum glycerol for the use in the phospholipid synthesis [9]. Zeuthen et al. reported that *Pf*AQP also facilitate ammonia diffusion, underscoring its importance in the release of ammonia produced from amino acid breakdown [12].

Targeted deletion of ortholog of *Pf*AQP in the rodent malaria parasite, *Plasmodium berghei* (PbAQP) resulted in delayed growth of parasite as compared to wild type [13]. Besides, PbAQP-null parasites were highly deficient in glycerol transport and mice infected with PbAQP-null parasites lived longer. These data indicated that PbAQP plays a significant role in the blood-stage development of the rodent malaria parasite and can be a probable candidate for development of anti-malarials [13]. Another report by Posfai et al. demonstrated that expression of host aquaporin ‘AQP3’ is considerably induced in *Plasmodium berghei* infected hepatocytes compared to uninfected cells [14]. Genetic disruption of AQP3 or treatment with its inhibitor ‘auphen’ resulted in reduced parasite load in liver and blood cells [14]. The same study also reported that AQP3-mediated nutrient transport plays a key role in parasite development [14].

In light of above facts highlighting the importance of *Pf*AQP, we looked for its inhibitor in the existing literature. A study by Chen et al. used *in silico* approach to show that erythritol, a permeant of *Pf*AQP itself has a deep ditch in its permeation passageway and may strongly inhibits *Pf*AQP’s functions of transporting water, glycerol, urea, ammonia, and ammonium [15]. This indicated the potential of this sweetener to act as a drug to kill the malarial parasite without causing side effects. Therefore, in the present study, we tested the effect of erythritol on different stages of malaria parasite through *in vivo* and *in vitro* experiments. Growth progression analysis suggests that erythritol treatment affects every asexual blood stage of parasite development. Activity against asexual blood stages of *Pf*3D7 and *Pf*RKL-9 was confirmed with values of IC_50_ (50% inhibitory concentration) to be 1.9 μM and 1.4 μM respectively. Our data demonstrate significant reduction in the percentage parasitaemia in erythritol treated mice infected with *P. berghei* ANKA. Combination of erythritol with known anti-malarial artesunate provide additive affect in inhibiting parasite growth both *in vivo* and *in vitro*. Moreover, erythritol showed anti-inflammatory properties owing to reduced proinflammatory cytokines in erythritol treated infected mice. Our cellular thermal shift assay clearly indicates the interaction of erythritol with *Pf*AQP. Also, we observed reduced amount of ammonia release across the parasite which indicates that erythritol is capable of blocking *Pf*AQP channel. Our complementation assays suggest that *Pf*AQP expression in yeast mutant restores its growth in hyperosmotic conditions and showed delayed growth in the presence of erythritol. Overall, our data strongly signify the ability of erythritol as a promising future drug molecule for malaria treatment.

## MATERIAL & METHODS

### *In-vitro* culture of *P. falciparum*

Laboratory strain of *P. falciparum* 3D7 and RKL-9 were cultured in RPMI 1640 (Invitrogen, USA) supplemented with 27.2 mg/L hypoxanthine (Sigma Aldrich, USA) and 0.5% Albumax I (Invitrogen, USA) using O+ erythrocytes in mixed gas environment (5% O2,5% CO2, and 90% N2). Parasites were synchronized by sorbitol selection of rings and percoll selection of schizonts.

### Parasite Growth Inhibition Assay (GIA)

A 72-h *in-vitro* growth inhibition assay was used to test the antimalarial activity of erythritol. The parasites were synchronized by treating with 5% sorbitol when ring stage was observed under microscope. 200 µL of the culture with parasitaemia of 1% rings and a haematocrit of 2% was added into a 96 well plate in triplicate and treated with erythritol (concentration range 250 nM to 50 µM). The antimalarial of the erythritol was estimated 72 h after treatment with it. Parasites of thin blood films stained with Giemsa were counted under a microscope. Total parasitaemia of treated cultures was compared with the control (only infected RBCs).

### Stage specific progression assays

To check the stage specific effect of erythritol on malaria parasite, *in-vitro* progression assays were performed at three distinct asexual stages of parasite. Briefly, synchronized cultures of ring (∼8-10 h), trophozoite (∼28-30 h) and schizont (∼36-40 h) stages of *Pf*3D7 strain were treated with 2 µM, 5 µM of erythritol in 96 well plate (1% parasitaemia; 2% haematocrit) for 72 hours. In other method, erythritol treatment was given at ring (∼8-10 h), trophozoite (∼28-30 h) and schizont (∼36-40 h) stages for 6 h followed by extensive washing to remove traces of drugs. Washed cultures were reseeded in 96 well plate (1% parasetemia; 2% haematocrit) for 72 h. Thin blood smears were prepared, stained with Giemsa stain and live parasites were counted under 100 x objective light microscope. Total parasitaemia of treated cultures was compared with the untreated one.

### Egress & Invasion assays

To assess the effect of erythritol on egress and invasion process of malaria parasite, schizonts stage culture (∼ 46 hpi; 5% parasetemia; 2% haematocrit) was treated with IC_50_, IC_90_ and IC_120_ concentration of erythritol for 10 h. Thin blood smears were prepared and stained with Giemsa. Schizonts and rings were counted under the 100x objective light microscope. Percentage egress was calculated as the fraction of schizonts ruptured in treatment and control during incubation time as compared with the initial number of schizonts at 0 h, using the formula as described below. Percent egress was then plotted considering the fraction of schizonts ruptured in control as 100% egress. To assess the effect of erythritol on parasite invasion, number of rings per schizont egress was calculated.

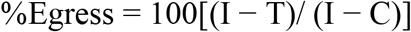

I, initial no. of schizonts; T, no. of schizonts in treatment; C, no. of schizonts in DMSO control.

### *In vitro* combination assays

Synchronized ring stage cultures of 3D7 and RKL-9 (1% parasetemia; 2 % haematocrit) were treated in combination with erythritol (0.5 µM, 1 µM, 2 µM, 4 µM and 8 µM) and artesunate (ART, 0.5 nM, 1 nM, 2 nM, 4 nM and 8 nM) in 96 well plate in fixed ratio method in triplicate. Both erythritol and artesunate were also subjected to individual testing. Thin blood smear slides were prepared followed by Giemsa staining. The untreated culture was taken as negative control. Around ∼2000 cells were counted under 100 x objective of light microscope. The percentage of parastemia was calculated as previously mentioned.

### *In vitro* growth inhibition assay for liver-stage parasites

Erythritol was added to 24-well culture plates (final DMSO concentration was 1%) containing human HepG2 cells which were seeded a day prior to the experiment and 0.5ml complete Dulbecco modified Eagle medium (DMEM) containing 10% fetal bovine serum (FBS), together with a cocktail of penicillin, streptomycin and amphotericin. Infection was initiated by adding 10,000 *Plasmodium berghei ANKA* sporozoites. Infected cultures were allowed to grow at 37^⟦^C in a 5% CO_2_ atmosphere for 48 h. Total RNA was extracted using TrizolTM (Invitrogen, USA) reagent and reverse transcription was performed using a cDNA synthesis kit (Applied biosystems). In a real-time PCR mix of 20 μl, a cDNA equivalent of μg RNA was used. The real-time PCR mix also contained *P. berghei* 18S rRNA specific primers.

### Gametocyte Exflagellation Assay

To initiate *P. falciparum* gametocyte cultures, synchronized asexual blood stage cultures were grown to a parasitaemia of 10-15%, treated with 2 µM erythritol containing medium for 4 days to remove asexual stages and maintained in complete RPMI/HEPES to allow gametocyte development. Stage V gametocytes were incubated with activation medium (100 nM xanthurenic acid (XA), 20% B+ human serum in RPMI1640/HEPES) in the presence of erythritol at 25^⟦^C. Activated gametocytes were spread on glass slides 5min after activation, fixed with methanol and stained with Giemsa (RAL Diagnostics, France). Around 200–250 gametocytes were scored to determine the percentage of rounded up gametocytes. The number of exflagellation centres was scored in gametocyte.

### *In vivo* combination assays

BALB c mice of six week age were divided into six groups with 5 mice in each one. Group 1 mice were left uninfected while malaria parasite *P. berghei* ANKA (1×10^6^) were passage intra peritoneally into each mouse of other five groups. Group 2 was treated with 100 mg/kg erythritol (Sigma-Aldrich, USA), Group 3 treated with 6 mg/kg artesunate, Group 4 treated with 60mg/kg artesunate, Group 5 treated in combination of erythritol (100 mg/kg) and artesunate (6mg/kg), and Group 6 was left as untreated (negative control). Drugs were orally administered to each mice once a day till twelth day post infection. Blood smears of the infected mice were stained with Giemsa. Parasitemia was determined by observing the stained smears under microscope. Survival of mice were monitored till 60 days post infection. The percentage of parastemia was calculated by using following formula:

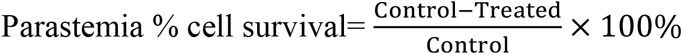

### Prophylactic and Suppressive assays

The prophylactic and suppressive efficacy of erythritol were investigated in six-week-old BALB/c female mice. The mice were distributed into 3 groups with 5 mice in each group. Group 1 was left untreated (positive control). For prophylactic experiments (group 2), oral administration of erythritol (100 mg/kg) was given four days prior to infection with malaria parasite and continued for another four days post infection. While in suppressive experiments (group 3) oral erythritol administration (100 mg/kg) was given from day 0 to eighth day of post infection. Mice were infected by intraperitoneal inocculation of 200 µl of blood diluted in PBS containing 1 × 10^6^ parasitized red blood cells, which was obtained from a donor mice. Blood from tail tip region of mice was taken on daily basis for slides preparation. Geimsa stained slides were examined under microscope and percentage of parastemia was determined by counting parasitized RBCs on atleast 4000 cells. Survival of mice was monitered for 20 days post infection.

### Evans Blue dye leakage assay

Evans Blue solution (2%) prepared in PBS was injected intraperitoneally (4 ml/kg of mice) in infected and uninfected mice. After 24 h of stain circulation, brain from one mouse from each group were removed under anaesthetized condition (by ketamine-xylazine cocktail, Sigma-Aldrich). Mice brains were placed in dimethyl formamide (Sigma-Aldrich) followed by incubation at 56°C overnight. Absorbance of the supernatant was measured at 610 nm in a microplate reader (Varioskan Flash, Thermo Fisher Scientific, USA). The dye leakage was calculated through extrapolation from the standard curve prepared for varying concentrations of evans blue.

### cDNA synthesis from mouse brain RNA and qRT PCR of pro-inflammatory cytokines

RT-PCR was performed to check the expression level of pro inflammatory cytokines in the brain cells of *P. berghei* infected mice treated with erythritol alone and in combination with artesunate. Excised mice brain tissue was dissolved in Trizol and stored in -80° C for RNA isolation. RNA from mice brain tissue was isolated by using RNA extraction kit (Thermo fisher) according to the manufacturer’s protocol. Total RNA was quantified and cDNA from isolated RNA was prepared by using HIGH-Capacity cDNA Synthesis KIT (Applied Biosystems) according to the manufacturer’s protocol. Each 10µL reaction mixture contain 1µl of cDNA, 0.5 µl of Fast STAR DNA SYBR Green reagents (Roche) and 0.5mM concentrations of primer. The PCR conditions consisted of an initial denaturation at 95 ° C for 10 min followed by amplification for 40 cycles of 10 s at 95°C, 5 s at 50°C and 20 s at 72°C with fluorescent acquisition at the end of each extension step. Amplification was immediately followed by a melt program consisting of 2 min at 60 ° C and a stepwise temperature increase of 0.2 ° C/s until 90°C with fluorescent acquisition at each temperature transition. GAPDH gene was used as a positive control.

### Molecular cloning, over-expression and purification of *Pf*AQP

Construct spanning 191 to 217 amino acids of PF3D7_1132800 (*Pf*AQP) was cloned in bacterial expression vector pET28a and expressed in BL21 (DE3) *E. coli* cells. Expression of *Pf*AQP was induced with 1 mM IPTG at 37°C for 3 h. Purification of recombinant protein was achieved by affinity chromatography.

### Raising polyclonal antisera against *Pf*AQP

Polyclonal antibodies were raised in house against *Pf*AQP in male BALB/c mice using purified recombinant protein as immunogens following standard protocol [16]. Briefly, animals were immunized with emulsion containing 1:1 ratio of freund’s complete adjuvant and antigen followed by three booster doses at an interval of 14 days. Booster doses comprise 1:1 ratio of freund’s incomplete adjuvant and antigen. Final bleed was collected and the end point titers of raised antisera were determined by western blotting.

### *In-vivo* expression analysis of *Pf*AQP

For *in vivo* expression analysis, *Pf*3D7 mixed stage asexual cultures (parasitemia ∼8%) were subjected to saponin lysis (0.15% w/v) followed by extensive washing of parasite pellet with 1x PBS to remove traces of haemoglobin. Protein extract of parasite lysate was resolved on SDS-PAGE and transfer to NC membrane followed by blocking in 5 % BSA for overnight at 4°C. Following blocking, the membrane was washed with PBS (Phosphate buffer saline) followed by probing with specific antisera against *Pf*AQP (1:500) for 2 h. Post washing with PBST (0.025%) and PBS, NC membrane was hybridized with horseradish peroxidise (HRP)-conjugated secondary antibodies for 2 h. Membrane was subsequently washed with PBST (0.025%) and PBS, and developed with DAB/H_2_O_2_ substrate.

### Immunofluroscence assays

Thin blood smears of mixed stage *Pf*3D7 cultures at 5% parasitaemia were fixed in methanol for 45 min at -20 °C, permeablized with 0.05% PBS/Tween 20, and blocked with 5% (w/v) BSA in PBS. Mouse anti-*Pf*AQP (1:200) was added as primary antibodies and incubated for 2 h at room temperature. Alexa Fluor 488 conjugated anti-mouse (1:500, green colour; Molecular Probes) was used as secondary antibodies. The parasite nuclei were counterstained with DAPI (40, 60-diamidino-2-phenylindole; Invitrogen, USA) and mounted with a coverslip. The slides were examined using a confocal microscope (Nikon Eclipse Ti2) with a 100x oil immersion objective.

### Cellular Thermal Shift Assay (CETSA)

*In vivo* interaction between erythritol and *Plasmodium* aquaporin was tested by Cellular Thermal Shift Assay (CETSA). Briefly, high parasitized *Pf*3D7 culture was pelleted down and subjected to RBC lysis using 0.15% saponin. Lysed pellet was extensively washed with PBS to remove traces of haemoglobin followed by RIPA lysis. Supernatant of lysate was separated out and treated with IC_50_ concentration of erythritol at four different temperatures (4° C, 40° C, 60° C and 80° C) for 5 minutes. Treated supernatant was resolved on 12% SDS-PAGE and transferred to nitrocellulose membrane. Blot was probed with anti-*Pf*AQP antisera and developed using ECL substrate. Anti-*Pf*GAPDH antisera was used to measure equal loading of samples. Recombinant *Pf*AQP was taken as positive control.

### *In-vivo* and *in vitro* ammonia detection

Brain samples of erythritol and artesunate treated mice were used for ammonia detection using Abcam-Ammonia Assay Kit (ab83360). Briefly, 10µgm of brain tissue was dissolved in ammonium chloride followed by centrifugation for 10 minutes. 50 of reaction mixture was added to standard and samples wells and 50 μL of background reaction mix to background control sample wells. Samples were incubated at 37°C for 60 minutes protected from light and the absorbance was recorded in a microplate reader at 570nm. Untreated mice were used as control.

Similarly, purified trophozoites of *Pf*3D7 were treated with IC_50_ and IC_90_ of erythritol. Parasites were washed with ice cold 1X PBS and then resuspended in assay Buffer. Culture media of erythritol treated parasite and untreated parasite were also used for ammonia detection. 50 of reaction mixture was added to each sample and was incubated at 37°C for 60 minutes protected from light. Absorbance was recorded in a microplate reader at 570nm.

### Functional complementation assays in yeast

Competent cells preparation and transformation of plasmids into *S. cerevisiae* were performed using the Frozen-EZ Transformation II TM protocol. Briefly, yeast cells were grown in YPD media (1% yeast extract, 2% peptone, 2 % glucose) and growth of transformants was made in selective medium (YNB containing 2% glucose) in the presence and absence of leucine for selection. For growth assays, cells were pre grown for 48 hours on YNB plates, resuspended in YNB till they reach OD _600_ of 0.2 and diluted in 10-fold dilution series. 5µL of each diluted cell suspension was spotted on agar plates supplemented with either 0.5M NaCl, 0.5M KCl, 1M Sorbitol or 1M erythritol and combination of erythritol with 0.5 M NaCl, 0.5 M KCl and 1 M Sorbitol. Growth was monitored for 2-4 days at 30 ° C.

## Results

### Erythritol inhibits *in vitro* growth of malaria parasites

The anti-malarial activity of erythritol on *Pf*3D7 and *Pf*RKL-9 strain was determined by growth inhibition assays (GIA). Our results showed that erythritol is effective against both strains and showed inhibitory effects in a concentration dependent manner as compared to untreated control. Half-maximal inhibition concentration (IC_50_) of erythritol against *Pf*3D7 and *Pf*RKL-9 were found to be 1.9 μM and 1.4 μM respectively (Fig. 1 A i, ii).

**Figure 1:**
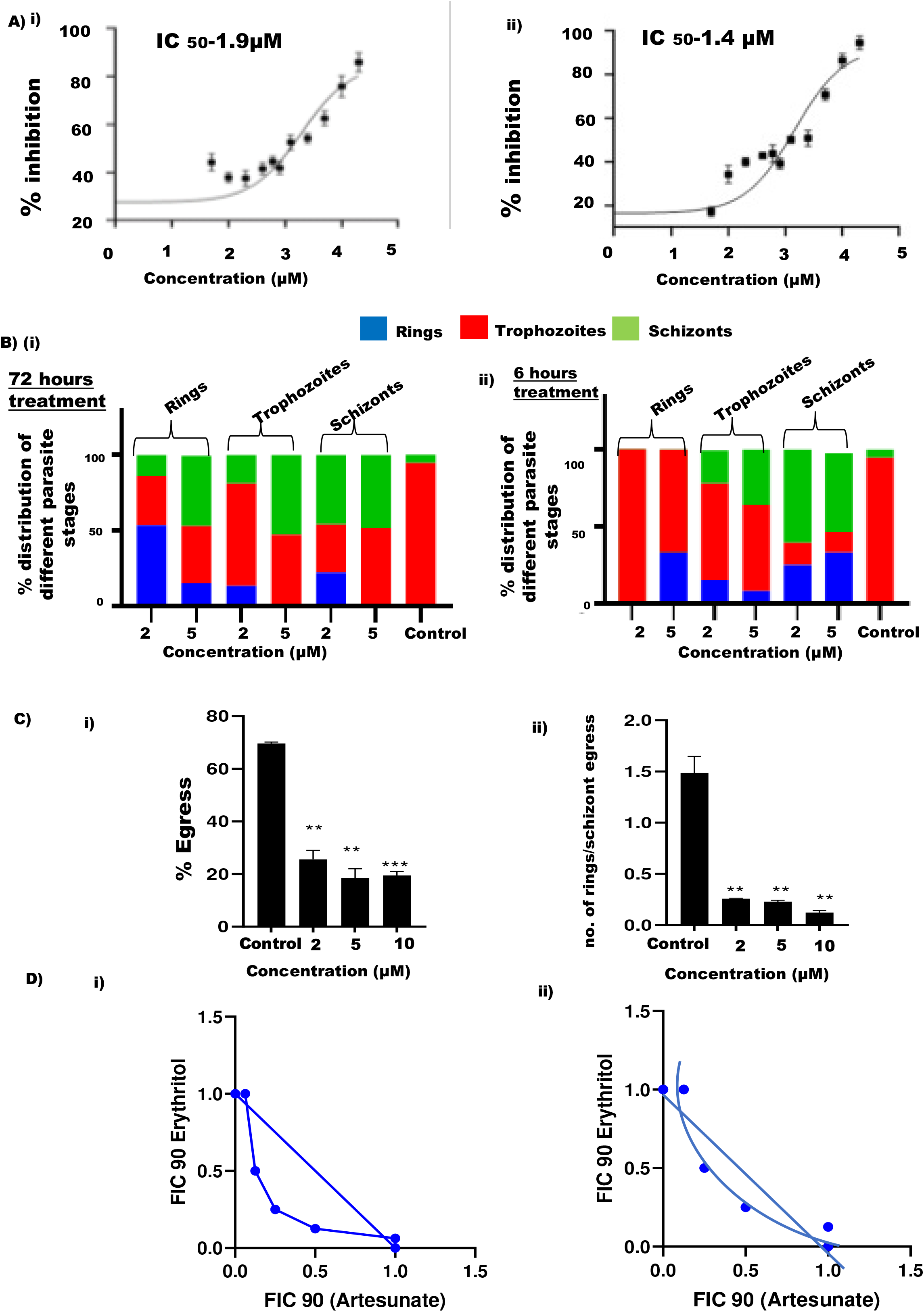
(A) Growth inhibitory effect of erythritol. The parasite culture synchronized at ring stage was treated with different concentrations (250 nM to 50 µM) of erythritol for 72 h, and the percent growth inhibition was estimated by Giemsa staining. IC_50_ values of erythritol in *Pf*3D7 (i) and *Pf*RKL-9 (ii) were evaluated by plotting growth inhibition values against the log concentration of erythritol. The experiment was done in triplicate, and the results were shown as mean values ± SD. **(B)** *Growth progression assays*. Graph shows percentage distribution of asexual stages of parasite following erythritol treatment. (i) Synchronized cultures of ring (∼8 -10 h), trophozoite (∼28-30 h) and schizont (∼36-40 h) stages of *Pf*3D7 were treated with IC_50_ and IC_90_ of erythritol for 72 hours. (ii) Erythritol treatment was given at different stages of parasite for 6 hours followed by extensive washing. Washed cultures were reseeded for 72 hours and parasites were counted using light microscope. Total parasitaemia of treated cultures was compared with the untreated control. **(C)** *Egress and invasion assays* (i) Late-stage schizonts were treated with different concentrations of erythritol. The relative inhibition in egress was determined by counting the number of remaining schizonts after 10 h of treatment as compared to control. (ii) Schizont stage *Pf*3D7 culture was treated with varying concentrations of erythritol. The ability to form rings per schizont egress was determined by counting the number of schizonts and rings after 10 h using giemsa stained smears. **(D)** *In vitro fixed ratio drug combination* assays. Isobologram showing synergistic effect of erythritol and artesunate on growth of *Pf*3D7(i) and *Pf*R539T(ii).

During the asexual phase of life cycle, the parasite undergo progression through different stages, i.e., rings, trophozoites, and schizonts. We investigated stage specific effect of erythritol on *Pf*3D7 using growth progression assays. Here, synchronized parasite culture of ring, trophozoite and schizont stage were treated with erythritol for 6 hours at their IC_50_ concentration and the effect of treatment was monitored till trophozoite stage of second cycle (72 h). In second set of experiment, erythritol treatment was continued to 72 h without drug removal. Untreated culture was used as a negative control. Growth progression defect was analysed by counting ∼3000 cells per giemsa stained slide in 100x objective of light microscope. Significant reduction of parasitemia at all three asexual stages of malaria parasite was observed in erythritol treated culture as compared to untreated control (Fig. B i, ii). In untreated culture, growth of the cells progressed normally. These data suggest that erythritol inhibit the growth of malaria parasite at all asexual stages.

We next analysed the effect of erythritol at three different concentrations (2 µM, 5 µM and 10 µM) in egress and invasion process of malaria parasite. Synchronized parasite culture of schizont stage (36-40 h) was treated with erythritol for 10 hours. Initial parasitaemia was around ∼ 1% for egress and invasion assays. Our data revealed that erythritol inhibited the egress of *Pf*3D7 in a dose dependent manner. Erythritol showed ∼70% egress inhibition at 2 µM while more than 80% inhibition at 5 µM and 10 µM concentration (Fig. 1C i). Number of rings formed per egress of schizont on various erythritol concentration were analysed and found to significantly decrease the number of rings formed in treated culture as compared to the untreated control (Fig. 1C ii).

Further, *in vitro* fixed ratio drug combination assays of erythritol with artesunate were performed to assess their combined effect on the growth of malaria parasite. Synchronized ring stage parasites of chloroquine-sensitive strain *Plasmodium falciparum* 3D7 & artemisinin-resistant strain *Plasmodium falciparum* R539T were treated with erythritol (0.5 µM to 8 µM) and artesunate (0.5 nM to 8 nM) at various concentrations for 72 h according to fixed ratio method. In addition, both drugs were tested individually. Untreated culture was used as negative control. We observed that percent parasitaemia inhibition was 94% when erythritol and artesunate were used together at their IC50 values. However, percent parasitaemia inhibition was 60% and 65% when erythritol and artesunate were used individually. These results clearly demonstrate that erythritol along with artesunate showed synergistic effect for killing the malarial parasite (Fig. 1D i, ii). Percent inhibition at different combinations of erythritol and artesunate are listed in table 1.

**Table 1:** Percent inhibition at different combinations of erythritol and artesunate.

| S.no | Combinations (artesunate + Erythritol) | % Inhibition in *Pf3D7* | % Inhibition in R539T |
| --- | --- | --- | --- |
| 1 | 0.5 nM + 8μM | 97% | 82.5% |
| 2 | 1 nM + 4μM | 92% | 83.5% |
| 3 | 2 nM + 2μM | 94% | 85% |
| 4 | 4 nM + 1μM | 95% | 82% |
| 5 | 8 nM + 0.5 μM | 96% | 89% |

### Erythritol treatment reduces *P. berghei* sporozoite load in HepG2 cells

To test the inhibitory effect of erythritol on sporozoite infection, we infected HepG2 cells with *P. berghei* sporozoites and treated with IC_50_ of erythritol for 72 h. The sporozoite infection was scored by real-time PCR analysis using *P. berghei* 18S rRNA specific primers. Our results clearly demonstrate that erythritol has an inhibitory effect toward *P. berghei* sporozoites (Fig. 2A). Erythritol showed no effect in the viability of HepG2 cells suggesting that it is specific toward sporozoites but not host cell (data not shown).

**Figure 2:**
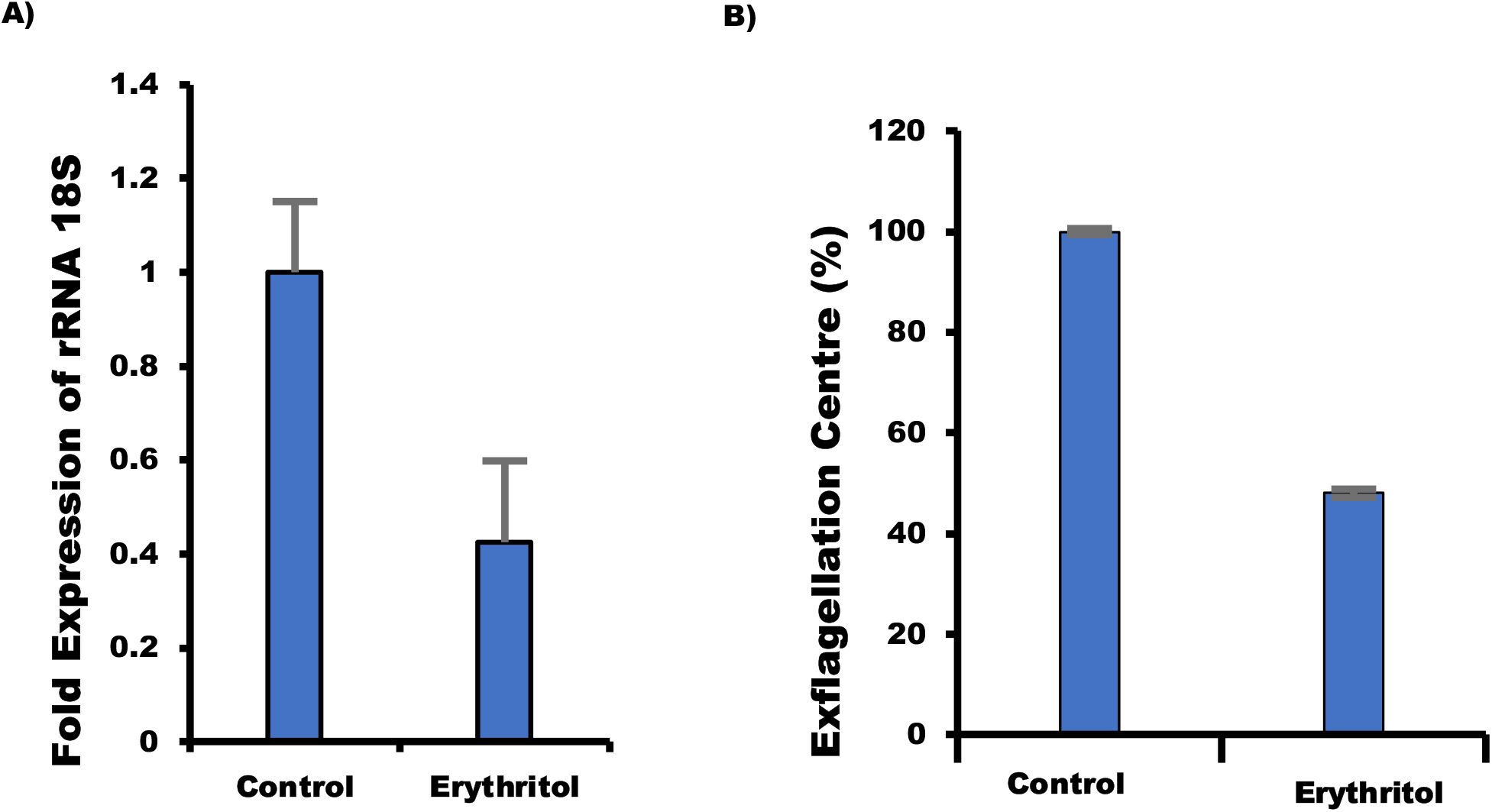
(A) Hep G2 Cytotoxicity Assay. HepG2 cells were infected with *P. berghei* sporozoites and treated with erythritol. Parasite growth was assessed via real-time PCR using *P. berghei* 18S rRNA specific primers. Graph shows fold expression of 18S rRNA in control and erythritol treated cells. **(B)** *Gametocyte Exflagellation Assay*: Treatment of male gametocytes with erythritol inhibits exflagellation. The number of exflagellation centers was scored and exflagellation efficiency of erythritol treated vs. untreated gametocytes is shown.

### Antimalarial action of erythritol against male gametogenesis

Erythritol was tested at its IC_50_ concentration in exflagellation assays. Mature gametocytes were preincubated with erythritol for 24 h before induction of gamete formation without removal of the compound. We observed that activated erythritol treated gametocytes did not form exflagellation centres, even when gametocytemia in the mouse blood was as high as 1% while around five exflagellation centres per microscopy field were present in control untreated activated gametocytes (Fig. 2B). Activated erythritol treated male gametocytes formed motile flagella, but they remained intracellular beating as a thick bundle of flagella, suggesting that they were unable to egress from the PV or the host erythrocyte. These data suggest that erythritol treatment blocks *P. berghei* gamete egress because of the inability to disrupt the membrane of the PV.

### Erythritol inhibited parasite load in mice

*In vivo* antimalarial efficacy of erythritol alone and in combination with artesunate were examined in *P. berghei* ANKA infected BALB/c mice. Mice were divided into six groups with five mice in each one. Group 1 mice were left uninfected while group 6 was not treated with any compound. Group 2, 3, 4 mice were treated with 100 mg/kg erythritol, 6 mg/kg artesunate, 60 mg/kg artesunate respectively while group 5 mice were treated in combination with 100 mg/kg erythritol and 6 mg/kg artesunate. Thin blood smears were made from each mice on daily basis up to twelve days. Our *in vivo* combination assay results demonstrated significant reduction of parasitemia in erythritol and artesunate treated group of mice as compared to untreated one (Fig. 3 A). On day 7, percentage of parasitemia was lowest in group 5 (∼5 %) that was treated in combination of erythritol and artesunate. Parasitemia in group 3, 4 and 5 was around 7% while in untreated control, it was ∼15 %. On day 12^th^ post infection, parasite load was reduced to ∼0.25% in erythritol treated group of mice as well as in those treated with combination of erythritol and artesunate (Fig. 3 A). Also, the survival rate of erythritol and artesunate treated animals, and those treated with combination of erythritol and artesunate were significantly greater than untreated control mice. Graph plotted as percentage survival versus time showed improvement in mortality rate in case of drugs treated mice (Fig. 3 B). Erythritol, artesunate (6 mg/kg), artesunate (60 mg/kg) treated mice survived up to 60, 12, 60 days respectively while those treated with and artesunate (6 mg/kg) and erythritol (100 mg/kg) both survived up to 60 days.

**Figure 3.**
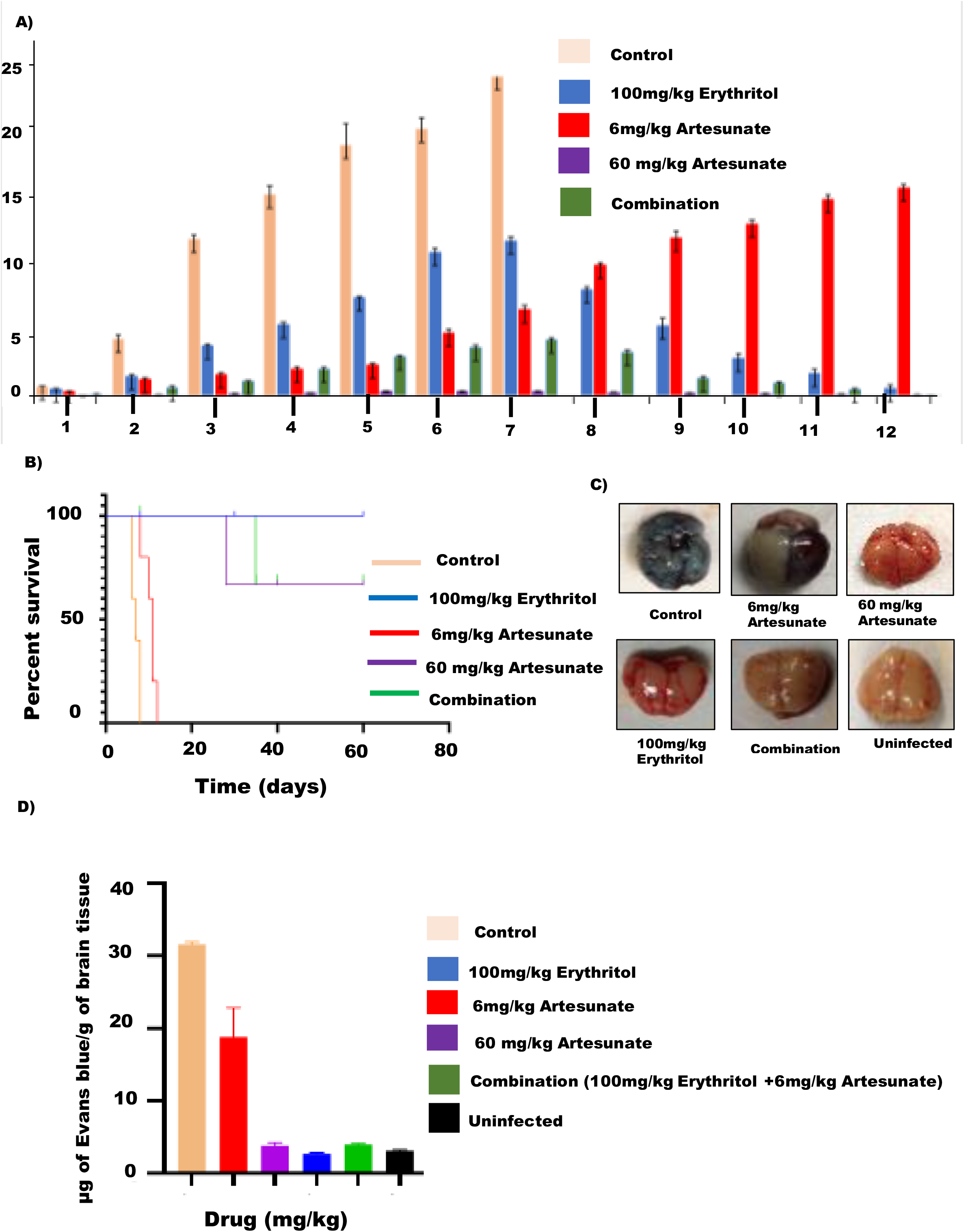
In vivo anti-malarial activity of erythritol. **(A)** Graph showing percent parasitemia in five experimental groups of mice (represented with different colors) treated with erythritol, artesunate and their combination. Significant reduction of parasitemia in erythritol and artesunate treated group of mice as well as in those treated with combination of both compounds was observed. **(B)** Graph showing percent survival of *P. berghei* infected mice treated with erythritol, artesunate and their combination. On day 7 all mice of untreated group died while more than 50% mice survived upto 60 days. **(C)** Comparison of blood–brain barrier damage in treated and untreated mice. Images of brains isolated from treated and untreated mice injected with evans blue displaying the color of dye in untreated mouse brain. **(D)** Graph showing amount of evans blue leakage due to damage in blood brain barrier in different groups of *P. berghei* infected mice (explained with different colors on graph).

We next performed evans blue leakage assay to determine the level of blood brain barrier damage caused by parasite in infected mice. The amount of evans leakage per gram of brain tissues were measured in all mice. Fig. 3 C showed leakage of evans blue dye in untreated mice, drug treated mice as well as in uninfected mice (positive control). Approximately 31.27 μg dye leakage per gram of brain tissue was observed in untreated mice, whereas it was 2.8 μg and 4.12 μg in erythritol treated mice and mice treated with combination of erythritol and artesunate respectively (Fig. 3 D). Evans leakage in mice treated with different doses of artesunate (60 mg/kg and 6 mg/kg) was 3.5 μg and 18.21 μg respectively. Overall, our data indicated negligible blood–brain barrier damage in treated mice as compared to untreated one.

### Erythritol exhibited significant prophylactic and suppressive activity against *Plasmodium berghei in vivo*

A prophylaxis is a treatment designed for prevention of a disease from occurring while, suppressive treatment reduces parasite load among infected model organism. We tested erythritol in prophylactic and suppressive treatment to compare its effect in *P. berghei* infected mice. In prophylactic treatment, mice were treated with erythritol 4 days prior to infection and continued for another four days post infection. While in suppressive treatment, oral administration of dose was given from day zero till eighth day post infection. The results from prophylactic study showed that erythritol delayed the onset of growth of malaria parasite in infected mice. Once erythritol administration stopped at day 4 post infection, a sharp increase in parasitemia was observed (Fig. 4A). While in suppressive assay, erythritol was able to significantly reduce the parasite load in infected mice as compared to untreated one (Fig. 4A). On day 8, parasitemia level was around 11% in both prophylactic and suppressive treated mice while it was 25% in untreated control mice. Graph plotted as percentage survival versus time also showed improvement in mortality rate in both prophylactic and suppressive treated mice (Fig. 4 B). These data indicate the potential of erythritol in prophylactic and suppressive treatment of malaria.

**Figure 4:**
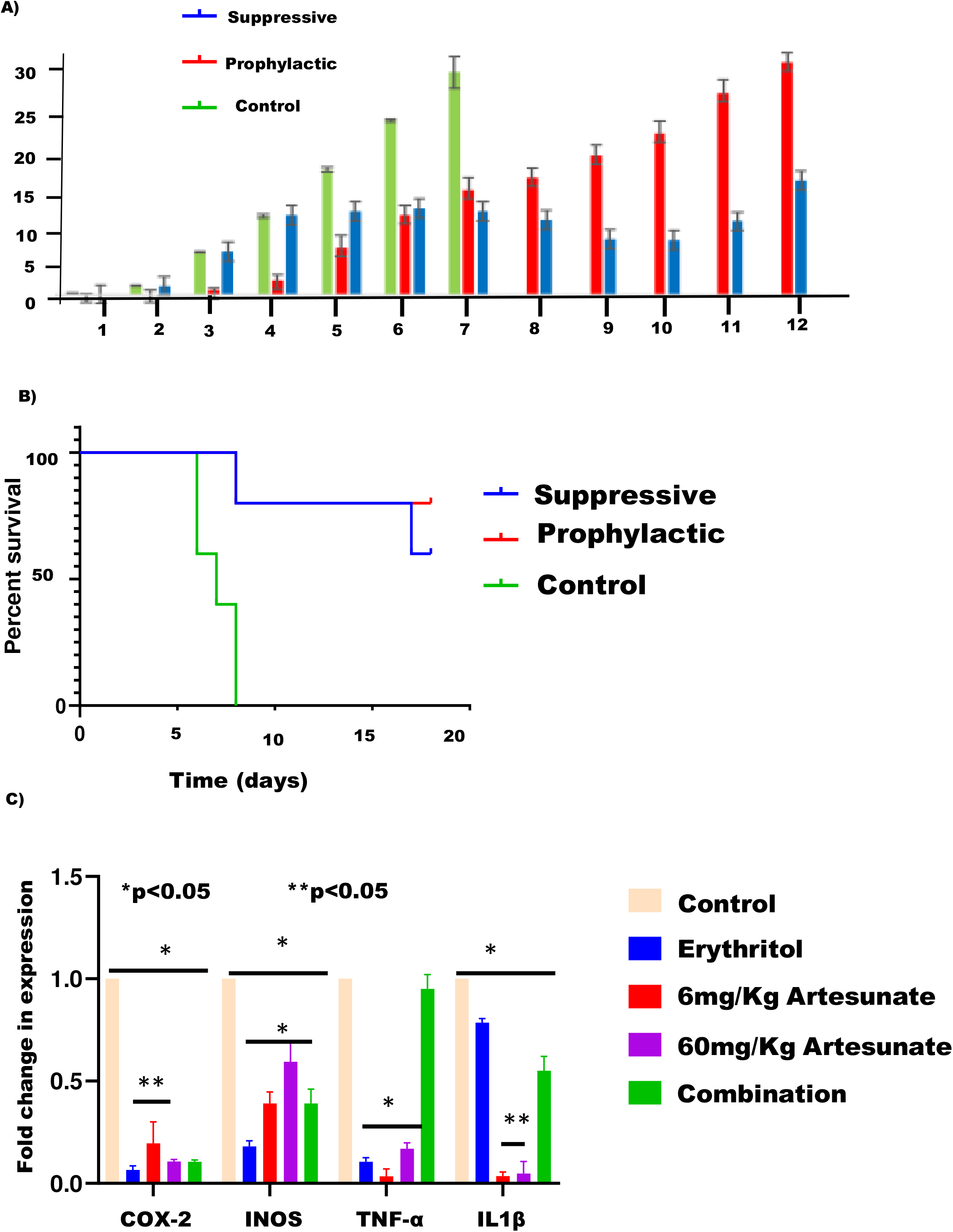
Prophylactic and Suppressive assays. **(A)** Graph showing percent parasitemia in three experimental groups of mice (represented with different colors) treated with erythritol. Significant reduction of parasitemia in prophylactic and suppressive treatment was observed. **(B)** Graph showing percent survival of *P. berghei* infected mice subjected with prophylactic and suppressive treatment with erythritol. **(C)** Level of pro-inflammatory cytokines in infected mice treated with erythritol, artesunate and their combination. Graphs showing transcriptional levels of the Cox2, INOS, TNF-α, IL1 β in mouse brain total RNA.

### Proinflammatory cytokines reduces upon erythritol treatment in *P. berghei* infected mice

Cytokines are key elements of the innate and adaptive immune responses eliciting inflammatory and febrile symptoms as well as activation of macrophages and play a key role in the pathogenesis of malaria [17]. We investigated the level of inflammation in the erythritol, artesunate and their combination treated infected mice by checking the expression pattern of Cox2 and pro-inflammatory cytokines INOS, TNFα, IL1β. Our results demonstrate a significant decrease in Cox2 (*P<0.05), INOS ((*P<0.05), TNF (*P<0.05) and IL1 β (**P<0.05) expression in erythritol, artesunate and their combination treated infected mice as compared to untreated control mice (Fig. 4C). These results suggest that erythritol has anti-parasitic and anti-inflammatory properties, which can inhibit the parasite growth and exert a direct effect on the host immune system.

### Cloning, expression and purification of *Plasmodium falciparum* aquaporin

With the aim of investigating the interaction of erythritol with *Pf*AQP, we cloned, expressed and purified *Pf*AQP. Construct spanning region from 191 to 217 amino acids of *Plasmodium falciparum* aquaporin (*Pf*AQP) was generated by cloning in pGEX-4T1 vector, followed by expression in *Escherichia coli* BL21 (DE3) cells. Figure 1A depicts a schematic representation of the domain organization of *Pf*AQP protein, highlighting the cloned region below the diagram (Fig. 5A). Expression of GST-fused *Pf*AQP was scaled up to purify protein from the soluble fraction using affinity chromatography (Fig. 5B i). Purified recombinant *Pf*AQP has molecular mass of 28.3 kDa. Identity of *Pf*AQP was confirmed by western blot using commercial anti-GST monoclonal antibodies (Fig. 5B ii). Polyclonal antisera against recombinant *Pf*AQP was raised in house in Balb/C mice. *In vivo* expression analysis on total parasite protein extract showed a single band of expected molecular weight for *Pf*AQP (Fig. 5C).

**Figure 5:**
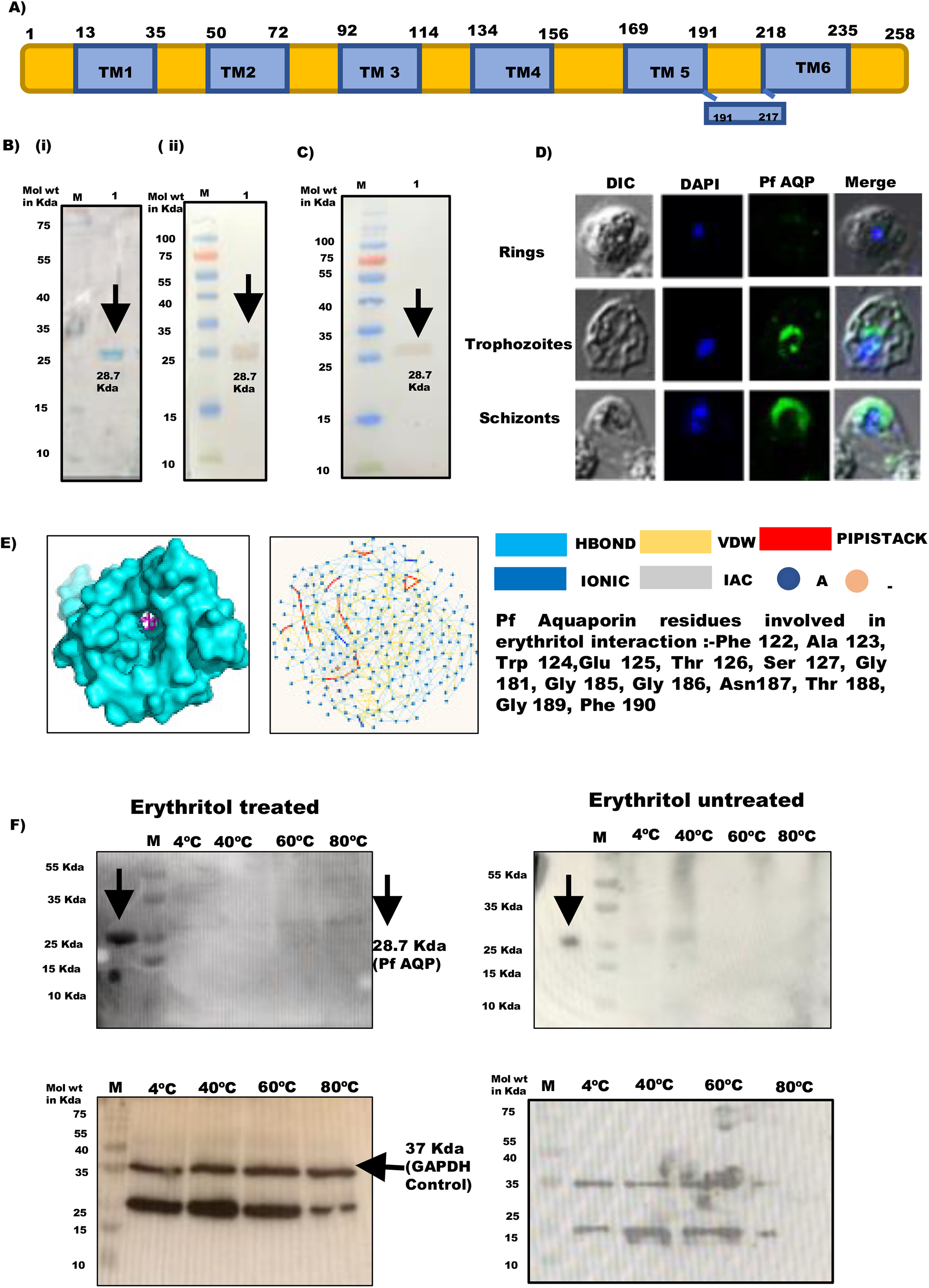
Expression of PfAQP and its binding with erythritol. **(A)** Schematic representation of *Pf*AQP gene showing cloned region below the diagram. **(Bi)** SDS PAGE of purified recombinant *Pf*AQP. Lane M: protein ladder; lane 1: purified recombinant *Pf*AQP. (ii) Western blot of purified recombinant *Pf*AQP using antihistidine antibodies. **(C)** Western blot analysis to test *in vivo* expression of *Pf*AQP in parasite lysate. Lane M: protein ladder; lane 1: parasite lysate. Blot was probed with anti-*Pf*AQP antibodies followed by secondary antibodies. **(D)** IFA images of *Pf*AQP at three different asexual blood stages of *Pf*3D7. Methanol fixed smear of *Pf*3D7 infected erythrocytes were stained with anti-*Pf*AQP antisera followed by incubation with alexa fluor conjugated secondary antibodies (Alexa Fluor 488, green colour). DIC: differential interference contrast image, DAPI: nuclear staining using 4^’^,6-diamidino-2-phenylindole (blue); *Pf*AQP: *Pf*PFD-6: mouse anti-*Pf*AQP antibodies (green); merge: overlay of *Pf*AQP with DAPI. (E) Left panel: Docked complex of *Pf*AQP with erythritol. Surface representation of *Pf*AQP is shown in cyan and erythritol is represented in stick and colored magenta. Right panel: Rin plot for erythritol interacting with *Pf*AQP. List of residues of *Pf*AQP interacting with erythritol are mentioned on the plot. **(F)** Interaction of *Pf*AQP with erythritol by CETSA. The drug-target engagement between the compound and recombinant protein was analyzed by subjecting the samples to thermal denaturation at 40, 60 and 80^⟦^C. Upper panel: Western blot analysis showing band of *Pf*AQP in erythritol treated (left panel) and untreated samples (right panel) at different temperatures (mentioned above the lanes). Lower panel: Western blot analysis showing band of GAPDH (loading control) in erythritol treated (left panel) and untreated samples (right panel) at different temperatures (mentioned above the lanes). Lane M: Molecular weight marker.

Immunofluorescence assays (IFAs) were performed to visualize the expression and localization of *Pf*AQP during the asexual stages of *Plasmodium falciparum* using protein-specific antibodies. Expression of *Pf*AQP begins at ring stage and continued to trophozoite and schizont stages (Fig. 5D). At early trophozoite and schizont stage, *Pf*AQP seems to surround the parasite plasma membrane (PPM), with very few foci observed in the parasite cytosol. However, to establish confinement of fluorescence at the PPM, costaining with PPM marker proteins would be required.

### Erythritol binds with *Plasmodium falciparum* aquaporin

Preliminary analysis of interaction was performed by docking solved structure of *Pf*AQP with erythritol. We observed that erythritol binds well into the cavity of *Pf*AQP with a binding energy of -64.01 KJ/mol (Fig. 5E i; left panel). To identify key residues involved in interactions, the residue interaction network (RIN) profiles of docked complex was generated using RING 2.0 web server (Fig. 5E i; right panel) [18]. RIN provides a visual interface to evaluate the stability of connections formed by amino acid residues at the contact sites [18]. Detail analysis of RIN plot suggested that Phe 122, Ala 123, Trp 124, Glu 125, Thr 126, Ser 127, Gly 181, Gly 185, Gly 186, Asn 187, Thr 188, Gly 189, Phe 190 of *Pf*AQP forms maximum number of interactions with erythritol (Fig. 1b).

We performed cellular thermal shift assay (CETSA) to evaluate the binding of erythritol to *Pf* Aquaporin. This technique involves the detection of target protein by monitoring the thermostability of native protein in the presence of its selective inhibitor [19]. Parasite lysate was treated with IC_50_ concentration of erythritol at four different temperatures (4° C, 40° C, 60° C and 80° C). Samples were resolved on 12% SDS-PAGE and transferred to nitrocellulose membrane followed by probing with anti-*Pf*AQP antisera. Analysis of band intensities demonstrated significant thermal protection of *Pf* aquaporin in the presence of erythritol, suggesting the binding of erythritol with *Pf* aquaporin (Fig. 5F). Anti-*Pf*GAPDH antisera was used to measure equal loading of samples. Recombinant *Pf*AQP was taken as positive control.

### Erythritol reduces the amount of ammonia release from parasite

Since *Pf*AQP is known to facilitate ammonia diffusion [12] and our CETSA results suggest that erythritol binds with *Pf*AQP, we studied the effect of erythritol on ammonia release by detecting the amount of ammonia in brain samples of erythritol and artesunate treated mice and also in mice where erythritol and artesunate were used in combination. We used Abcam-Ammonia Assay Kit which is based on the colorimetric assay where ammonia is converted to a product that reacts with the OxiRed probe to generate color that can be quantified by using a spectrophotometric plate reader. Our results clearly demonstrate that a significantly high amount of ammonia was present in untreated mice brain as compared to erythritol and artesunate and their combination treated mice (Fig. 6A). The amount of ammonia presents in erythritol and artesunate treated mice brain was similar to uninfected mice. These results suggest that erythritol reduces the parasite load in infected mice and, binds with *Pf*AQP, subsequently reducing the amount of ammonia release in the host tissue.

**Figure 6:**
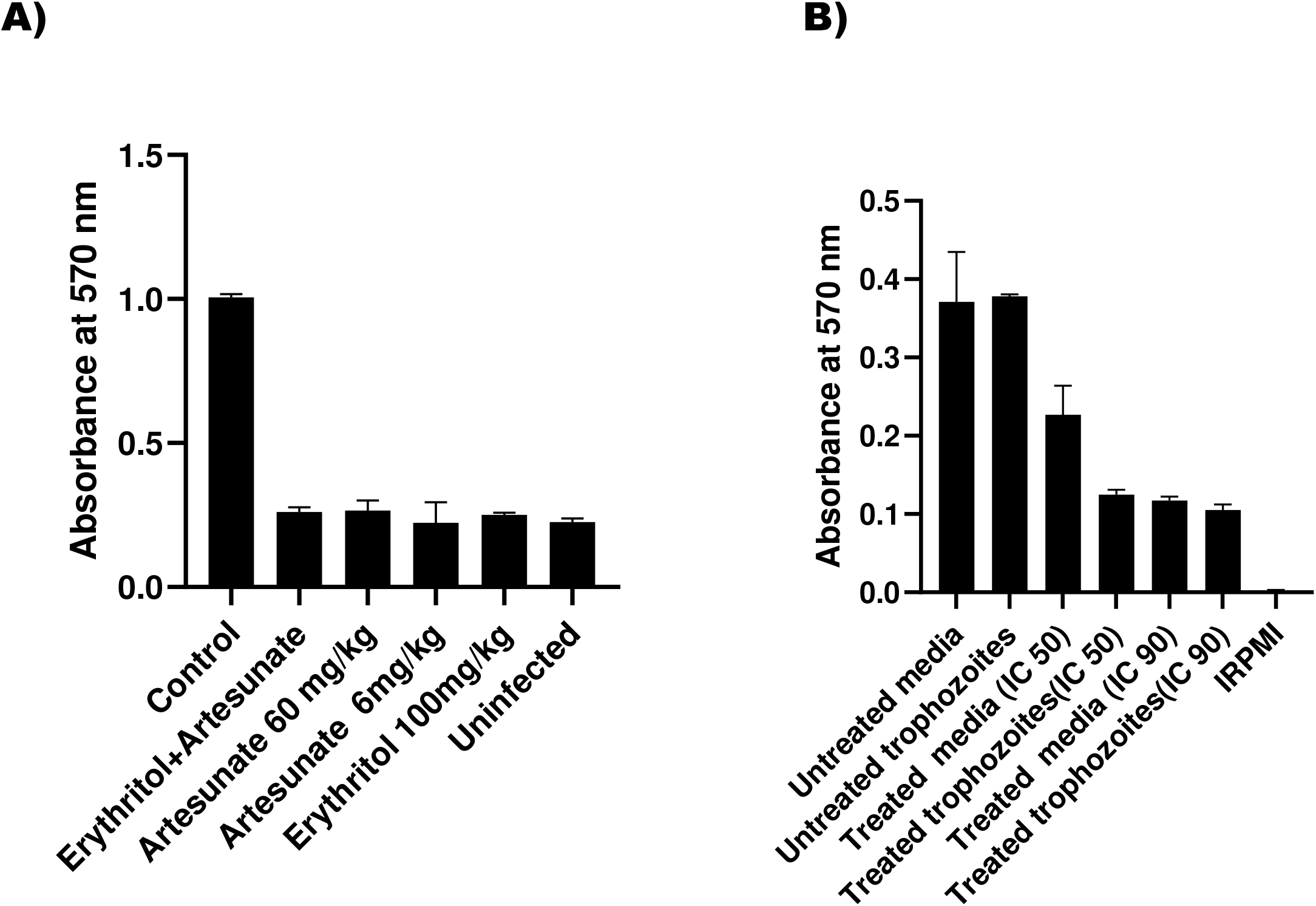
*In-vivo* and *in vitro* ammonia detection in erythritol treated parasite. **(A)** Amount of ammonia in brain samples of erythritol, artesunate and their combination treated mice (as labelled below the bars) was quantified by measuring the absorbance at 570nm using Abcam-Ammonia Assay Kit. Error bars represent standard deviation among three replicates. **(B)** Amount of ammonia was quantified in erythritol, artesunate and their combination treated treated parasite and their culture media (as labelled below the bars) by measuring the absorbance at 570nm. Error bars represent standard deviation among three replicates.

Similarly, purified trophozoites of *Pf*3D7 were treated with IC_50_ and IC_90_ of erythritol and amount of ammonia was quantified in treated parasite and its culture media. Our results show that the amount of ammonia is significantly less in erythritol treated parasite and its culture media as compared to untreated parasite and its culture media (Fig. 6B). Ammonia was quantified in erythritol dissolving solution IRPMI as a negative control. These results further suggest that erythritol treatment kills the parasite and blocks *Pf*AQP thereby reducing the amount of ammonia release in culture media.

### Erythritol selectively inhibits *Pf*AQP and *Pf*AQP expression restores growth of mutant yeast cells

We performed osmosensitivity assays in yeast to study the function of *Pf*AQP during the hyperosmotic condition. We used wild type yeast strain BY 4742 having the ability to excrete glycerol and its mutant BY 4742 ΔFsp-1 which is unable to excrete glycerol out of the cell. Similarly, wild type yeast strain YSH 202 and its mutants YSH 690 GPD1Δ, YSH 642 GPD 1Δ GPD 2Δ were also used in complementation experiments which are unable to synthesize glycerol. These mutant yeast strains were not able to grow in the presence of hyperosmotic conditions. Mutant yeast strains BY 4742 ΔFsp-1, YSH 690 GPD1Δ and YSH 642 GPD 1Δ GPD 2Δ were transformed with the plasmid mediating expression of codon optimized *Plasmodium falciparum aquaporin* (pYX242_*Pf*AQPcod-opt), *Plasmodium falciparum aquaporin* (*Pf*AQP; pYX242_PfAQP) and human AQP9 (pYX242_ Human AQP9). Before transformation, the phenotype of the BY4742 wild-type, YSH 202 wild type and their mutant strains were analysed on YP-agar plates containing high salts namely 0.5 M Nacl, 0.5 M Kcl, 1 M sorbitol and 1 M erythritol. Our results show that growth of BY 4742 ΔFsp-1, YSH 690 GPD1Δ and YSH 642 GPD 1Δ GPD 2Δ mutants were inhibited while BY4742 and YSH 202 wild type strain grew normally on hyperosmotic environment (Fig.7 A, B, C). To analyze phenotype of transformed cells in hyperosmotic conditions, cells were pre-grown in liquid medium till O.D reached to 0.2 and then serial diluted cells (1:10, 1: 100, 1:1000 and 1:10000) were spotted onto YPD plates. The BY 4742 ΔFsp-1 mutant transformed with codon optimised *Pf*AQP and human AQP9 restore its growth almost like wild type in the presence of high salt concentration whereas *Pf*AQP showed reduced growth phenotype under same conditions. Also, growth phenotype of codon optimised *Pf*AQP was decreased in YPD medium supplemented with 1M erythritol alone or in combination with other salts (Fig. 7A). YSH 690 GPD1Δ transformed cells restore growth to some extent (Fig. 7 B) while YSH 642 GPD 1Δ GPD 2Δ transformed cells did not confer any obvious growth phenotype on hyperosmotic environment (Fig. 7 C). These results suggest that expression of codon optimized *Pf*AQP in ΔFsp-1 mutant functionally complement its growth in the presence of hyperosmotic conditions. However, ΔFsp-1 mutant transformed with codon optimized *Pf*AQP showed delayed growth in the presence of erythritol. These data further suggest that erythritol behaves as an inhibitor of *Pf* aquaporin that blocks its channel and function of transporting solutes across the cell.

**Figure 7:**
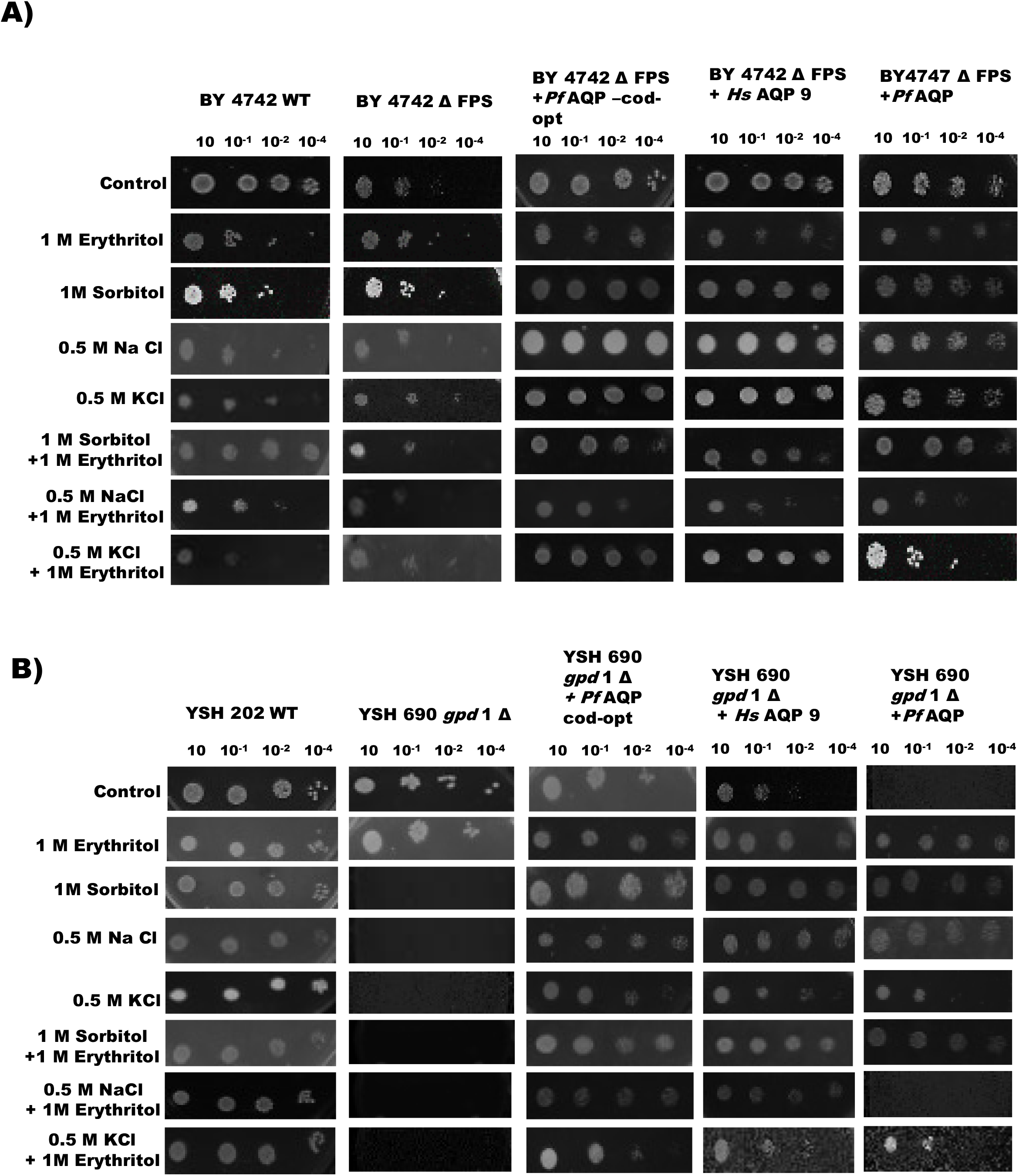
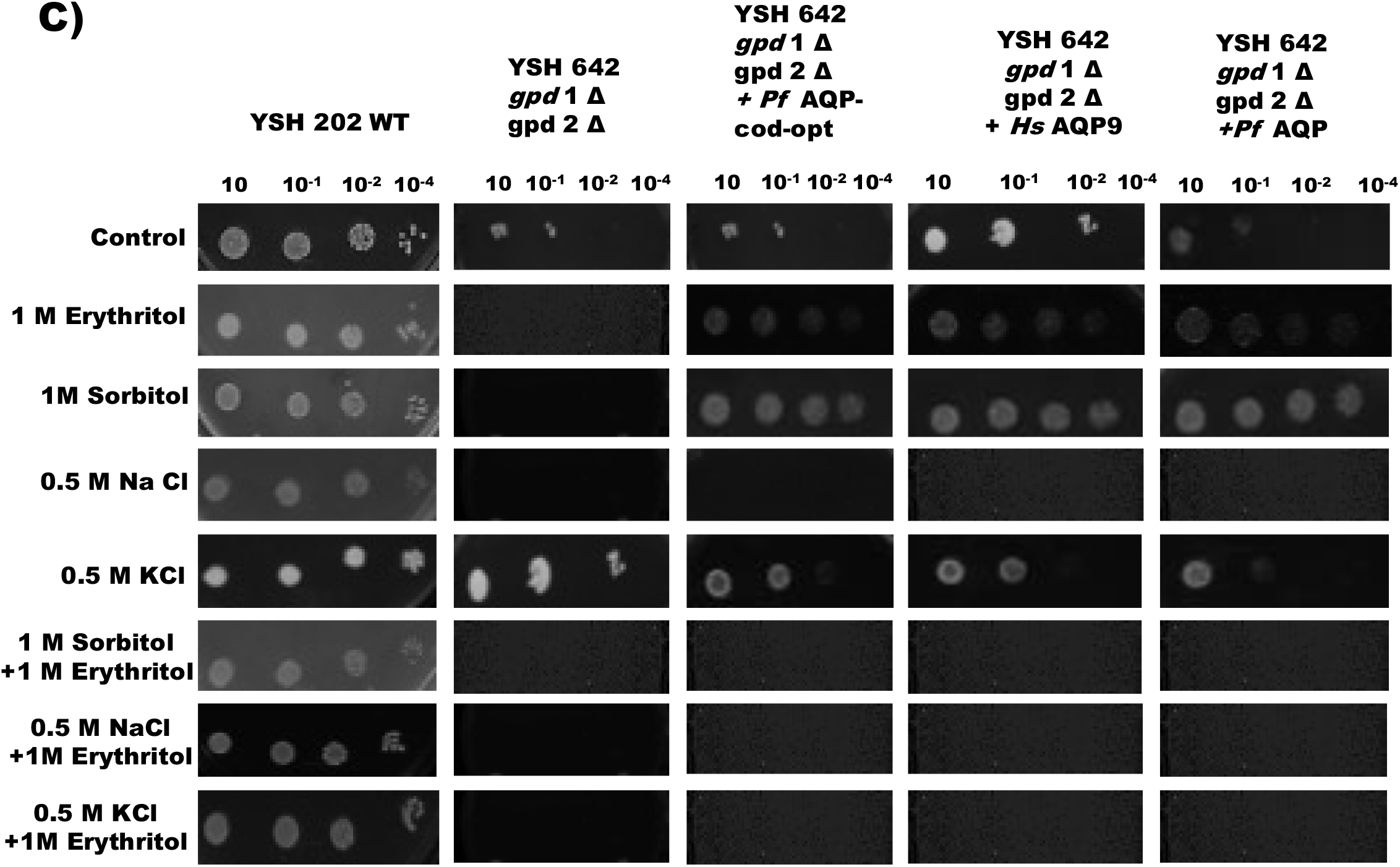
Functional complementation assays in yeast. Phenotype analysis of yeast cells transformed with *Pf*AQPcod-opt, human AQP9 and *Pf*AQP (labelled above the spots) in BY 4742 ΔFsp-1 **(A)**, YSH 690 GPD1Δ (B) and YSH 642 GPD 1Δ GPD 2Δ (C) in hyperosmotic conditions (as labelled on the figure). Cells were pre-grown in liquid medium till O.D reached to 0.2 and then serial diluted cells (1:10, 1: 100, 1:1000 and 1:10000; as labelled above the spots) were spotted onto YPD plates.

## Discussion

*Plasmodium falciparum* (*Pf*) is the major cause of severe malaria and is responsible for high mortality rates associated with the disease [20]. The ability of intraerythrocytic *Pf* to interact with endothelial receptors mediates the process of cytoadherence which is the key cause for the pathology of the disease [21]. Cerebral malaria is the most severe form of *Pf* infection and its pathogenesis is the result of cytoadherence and sequestration of iRBCs in brain microvasculature that often leads to impaired consciousness. Coma is the severe manifestation, and is regarded as clinical hallmark of the disease [20]. Although prophylaxis and chemotherapeutics has led to decrease in global mortality rate, yet developing resistance of malaria parasite to available drugs raises concerns about control of the disease. Lack of a successful vaccine for malaria signifies the constant need for identification of novel drug targets and their anti-malarials.

A previous study demonstrated that erythritol, a permeant of *Pf*AQP has a deep ditch in its permeation passageway and may inhibit *Pf*AQP’s functions of transporting various solutes [15]. This data provided a clue for the potential of this compound to act as a drug for malaria parasite. In light of the above report, we tested the anti-malarial activity of erythritol on different stages of malaria parasite through *in vivo* and *in vitro* experiments. We performed growth inhibition assays which clearly demonstrate that erythritol inhibited growth of blood stage *P. falciparum* 3D7 with an IC_50_ value of 1.9 μM. It was also almost equally active against chloroquine drug-resistant strain *Pf*RKL-9 (IC_50_:1.4 μM). Stage specific studies using synchronized cultures showed that erythritol prevented growth of all asexual blood stages of parasite including ring, trophozoite and schizonts with rapid killing. Further, our egress and invasion assays depict that the schizont stage parasites treated with erythritol could not egress and re-invade the fresh erythrocyte. Owing to the fact that anti-malarials show resistance with time, they are nowadays developed as combination therapies which are known to reduce the development of drug resistance and further improve efficacy of drugs [22]. Therefore, we assessed the effect of erythritol in combination with a known clinically approved anti-malarial artesunate on growth of parasite *in vitro*. Our data showed that both compounds have pronounced effect in inhibiting parasite growth when tested together. The combination of erythritol with artesunate in a fixed ratio method strengthen its effect on growth inhibition by showing synergistic effect in killing the malarial parasite. These results provide evidence for the chemopreventive potential of using erythritol in malaria infections that can be enhanced by combination with conventional antimalarials. This is the first report showing the *in vitro* anti-plasmodial activity of erythritol which is a 4-carbon sugar alcohol, often used as a sugar substitute and food additive [23]. Erythritol is almost non-caloric and is safe for diabetics as it does not affect blood glucose levels [23]. These properties prompted us to explore erythritol further for its binding with *Pf*AQP and as anti-malarial agent.

Since aquaglyceroporin play a role during multiple stages of the parasite cycle [13], we studied cross-stage inhibitory potential of erythritol. Malaria infection in humans starts after injection of sporozoites by anopheline mosquitoes, where the parasites invade and multiply in the liver [24]. We explored the effect of erythritol in blocking sporozoite infection *in vitro*. In the compound-treated HepG2 cells, the parasite burden was significantly reduced as compared to that for control. These data clearly demonstrate that apart from asexual blood stages, erythritol is also effective at sporozoites in preventing infection in liver cells. We also identified erythritol to act upon the mature gametocytes directly to inhibit processes unique to gamete formation. *P. berghei* gamete formation and host cell egress is readily observable *in vitro*. While a female gametocyte forms a single macrogamete, the male gamete undergoes three mitotic divisions and produces eight flagellated microgametes in just 10–15 min [25]. Exflagellation occurs when male microgametes use microtubule-based movements to leave the erythrocyte host and bind egressed macrogametes. These exflagellation centers made of gametes attached to erythrocytes, can be readily observed by microscopy. Our results indicate that the egress efficiency of male gametocytes, as revealed by counting exflagellation centers, was markedly reduced following erythritol treatment. This complementary activity of erythritol on the sexual stages of the parasites suggests its potential to reduce malaria transmission and its action on the sporozoite stage indicates its plausible role in chemoprevention. Overall, these results demonstrate antimalarial activity of erythritol against liver sporozoite stages, gametocytes and asexual blood stages of parasite. Several anti-malarial agents that are effective against multiple life-cycle stages have been reported in the past [26, 27]. Baragaña et al. described DDD107498, a compound that showed antimalarial activity against asexual blood stages, liver sporozoite stages as well as gametocyte stages of parasite [26].

We next evaluated *in vivo* antimalarial activity of erythritol alone and in combination with artesunate in mouse malaria model of *Plasmodium*. Our results indicated significant parasitaemia suppression in infected mice that were administered with erythritol alone and also in combination with artesunate. Percentage of parasitaemia was lowest in infected mice that was treated in combination of erythritol and artesunate. Also, erythritol prolonged the mean survival time of the mice indicating that it suppressed *P. berghei* and reduced the overall pathologic effect of the parasite on the mice. We next tested erythritol in prophylactic and suppressive treatment in *P. berghei* infected mice. Prophylactic treatment reduced the total burden of *P. berghei* infection and substantially lessened disease severity in mice. These findings highlight the potential of erythritol in prophylactic regimen for malaria chemoprevention. Also, our data for suppressive treatment indicated that erythritol has significant anti-malarial activity against *P. berghei* in mice evidenced by the percentage of parasitaemia inhibition.

Our *in vivo* studies also demonstrated that erythritol has cytokine-modulating effect apart from anti-malarial activity. Following erythritol treatment, there is significant reduction in the levels of pro-inflammatory cytokines in *P. berghei* infected animals. The dual anti-malarial and cytokine-modulating effects of erythritol render it a potential adjunctive therapeutic. Additionally, it is tempting to speculate that the increase in mean survival time of infected mice observed in the present study may be attributed to erythritol mitigating the cytokine storm.

With the aim of studying the interaction of erythritol with *Plasmodium falciparum* aquaporin, we cloned a construct spanning region from 191 to 217 amino acids of *Pf*AQP using pGEX-4T1 vector and overexpressed in soluble form in a bacterial expression system [BL21 (DE3) *E. coli* cells]. *In vivo* expression is clearly evident from our western blot analysis on total parasite protein extract which shows a single band of expected molecular size for *Pf*AQP. Our immunofluorescence data clearly depict that expression of *Pf*AQP begins at ring stage and continued to trophozoite and schizont stages. We explored the interaction of erythritol with *Pf*AQP by docking solved structure of *Pf*AQP with erythritol. Our results depict that erythritol interacts with *Pf*AQP with a binding energy of -64.01 KJ/mol. To confirm this interaction, we performed cellular thermal shift assay (CETSA) where parasite lysate was treated with IC_50_ concentration of erythritol at four different temperatures. Our data indicate significant thermal protection of *Pf*AQP in the presence of erythritol which is indicative of binding of erythritol with *Pf* aquaporin. This interaction is likely to block the *Pf*AQP channel for transporting solutes across the cell and provide it anti-parasitic property.

A study by Zeuthen et al. reported that *Pf*AQP assist ammonia diffusion, which is produced from amino acid breakdown [12]. We quantified the ammonia in brain samples of erythritol and artesunate treated mice as well as in erythritol treated trophozoites of *Pf*3D7 to study the effect of erythritol on ammonia release. Our results suggest that erythritol treatment reduce the amount of ammonia release in the infected mice brain and also in culture media of treated parasite. These findings clearly demonstrate that erythritol is capable of blocking *Pf*AQP channel thereby reducing the amount of ammonia release across the parasite.

We next performed complementation assays in yeast to study the function of *Pf*AQP during the hyperosmotic conditions. Yeast cells accumulates glycerol as an osmo-protectant when growing under hyper osmotic conditions [28, 29]. Fps 1 is the osmogated glycerol export channel in *S. cerevisiae* that play a central role in osmoregulation and maintain glycerol levels inside the cell [28, 29]. Mutation of the yeast Fps gene affects glycerol excretion in yeast cells, as compared to the wild-type strain. When we transformed yeast BY 4742 ΔFsp-1 mutant cells with codon optimized *Pf*AQP, they showed wild-type growth phenotype under hyperosmotic conditions. These data clearly suggest that *Pf*AQP complements the function of transporting solutes (glycerol) in yeast. However, BY 4742 ΔFsp-1 mutants transformed with codon optimized *Pf*AQP showed delayed growth in the YPD medium supplemented with erythritol. This indicate that erythritol is able to block aquaporin channel under hyperosmotic conditions. Our data supports a previous report where bioinformatics study had indicated the capability of erythritol in inhibiting *Pf*AQP function of transporting water and solutes across *P. falciparum* plasma membrane [15].

Overall, our data suggest that erythritol has anti-plasmodial and anti-inflammatory activity for malaria and when combined with current anti-malarials, it may be useful as a new and more effective preventive and therapeutic agent against the disease. Erythritol has potent activity profile against multiple life-cycle stages, a novel mode of action and excellent drug-like properties which has major implications for ensuring patient compliance and practical deployment. Its complementary activity on the sexual stages of the parasites makes it more potential to reduce malaria transmission. Since erythritol is a sweetener generally considered safe, it can prove to be an effective compound that kill the malarial parasite without causing serious side effects. We suggest that clinical trials can be performed to observe the effect of erythritol along with known antimalarials.

## Acknowledgements

**JK** is DST/INSPIRE fellow (Reference number: 2017/IF70636). **VK** is supported by Research Associateship Program of Department of Biotechnology, Govt. of India. **AB** is supported by National Post-doctoral fellowship, SERB, India (Fellowship reference no. PDF/2019/000334). **RKS** is CSIR-SRF, **SG** is senior research associate ship under the scientists’ pool scheme (Reference number: 13(9161-A)/2021. **SP** is grateful for the funding support from Shiv Nadar Foundation. Funding from Intensification of Research in High Priority Areas (IRHPA) of Science and Engineering Research Board (SERB) and the National Bioscience Award from DBT for SS is acknowledged.

Strains YSH642, YSH690 and YSH202-w303-1A wt and plasmids namely pYX242_Human AQP9, pYX242_*Pf*AQP and pYX242_*Pf*AQP-opt have been kindly gifted by Prof. Kristina Hedfalk; Department of Chemistry & Molecular Biology, University of Gothenburg, Sweden. Strains BY4742 and BY4742 fpsΔ have been kindly gifted by Prof. Eric Beitz; Department of Pharmaceutical and Medicinal Chemistry, Pharmaceutical Institute, University of Kiel, Germany.

## Declaration of interests

Authors declare no conflict of interests.

## Author contributions

**JK** conducted experiments, analyse data and helped in manuscript preparation. **VK** Experimental design, experimentation, data analysis and manuscript preparation. **AB**: Experimental design, experimentation, data analysis, manuscript writing. **RKS** conducted liver stage and gametocyte stage experiments. **SG** helped in experimental design and data analysis. **SP** helped in experimental design and data analysis. **KS, JB, NM** helped in data analysis and manuscript editing. **SS** conception of idea, experimental design, data analysis, manuscript writing and approval of final draft.

